# The kinase domain of RIPK3 tunes its scaffolding functions

**DOI:** 10.1101/2025.04.29.651198

**Authors:** Shene Chiou, Komal M. Patel, Adele Preaudet, James A. Rickard, Christopher R. Horne, Samuel N. Young, Sarah E. Garnish, Anne Hempel, Cathrine Hall, Joanne M. Hildebrand, Andrew J. Kueh, Tracy L. Putoczki, Edwin D. Hawkins, Andre L. Samson, James M. Murphy

## Abstract

The pro-inflammatory programmed cell death pathway, necroptosis, relies on phosphorylation of the terminal effector, MLKL, by RIPK3. RIPK3-deficient mice or those harboring the kinase-inactivating mutation, RIPK3^K51A^, are ostensibly normal in the absence of challenge, indicating that RIPK3 and its kinase activity are dispensable for development. However, another kinase-inactivating mutation, RIPK3^D161N^, results in embryonic lethality in mice due to widespread apoptosis. As a result, the RIPK3^D161N^ mutation is thought to confer a toxic gain-of-function. Here, to further explore the impacts of RIPK3 inactivation, we compared the stability and cellular interactions of RIPK3^D161N^ and RIPK3^K51A^ to a third previously-uncharacterized kinase-dead variant, RIPK3^D143N^. Here, we show that RIPK3^K51A^ was unstable and did not associate with RIPK1, RIPK3^D161N^ was unstable but interacted with RIPK1, whereas RIPK3^D143N^ was stable and bound RIPK1 in a manner comparable to wild-type RIPK3. Thus, all three variants scaffold differently, suggesting that the assembly of cell death machinery by RIPK3 is finely tuned, not just by its kinase activity, but also by the conformation of its kinase domain. Physiologically, *Ripk3^D143N/D143N^* mice exhibited a partially penetrant lethality *in utero*. However, once born, *Ripk3^D143N/D143N^* mice were fertile and phenotypically indistinguishable from wild-type mice in the absence of challenge, with RIPK3^D143N^ mutation protecting mice lacking intestinal epithelial Caspase-8 from necroptotic ileitis. Our studies support the idea that RIPK3 is a nexus between apoptotic and necroptotic signaling, and highlight the importance of considering kinase domain conformation in RIPK3 inhibitor development.

## Introduction

Necroptosis is a caspase-independent, pro-inflammatory programmed cell death modality that has been implicated in the pathology of a wide range of diseases. Lysis caused by necroptotic cell death releases damage-associated molecular patterns (DAMPs) into the extracellular milieu^1^. DAMP release, in turn, provokes an immune response, which reflects the likely origins of the pathway as an altruistic cell death mode that combats pathogens^2, 3, 4, 5, 6, 7, 8, 9^. Over the past decade, intense interest has centred on the role of dysregulated necroptosis in disease pathologies, particularly in ischemic injury and inflammatory diseases in the kidney, lung, gut and skin^10, 11, 12,13, 14, 15, 16, 17, 18, 19, 20, 21^.

Necroptotic signaling is initiated by ligation of death receptors, such as TNF Receptor 1 by TNF, or pathogen receptors, such as Toll-like receptors and the intracellular sensor protein, ZBP1, by microbial ligands. Under specific contexts, for example when the cellular inhibitors of apoptosis proteins (cIAP) E3 Ubiquitin ligase family and pro-apoptotic caspases are depleted, necroptosis can ensue by prompting assembly of a cytosolic signaling platform known as the necrosome. This megadalton complex is nucleated by the Receptor-interacting serine/threonine protein kinases (RIPKs), RIPK1 and RIPK3^22, 23^. RIPK1 and RIPK3 assemble into hetero-oligomers through the functional amyloid-forming RIP homotypic interaction motif (RHIM) within the sequence C-terminal to each kinase domain^24, 25, 26^. Autophosphorylation of both RIPK1 and RIPK3 is essential for necroptotic signaling, with autophosphorylation within the RIPK3 kinase domain C-lobe (T224/S227 in human RIPK3^27, 28, 29^; T231/S232 in mouse RIPK3^30^) essential for facilitating binding to, and recruitment of, the terminal pathway effector, the MLKL (mixed lineage kinase domain-like) pseudokinase, to the necrosome^29, 31^. Subsequent phosphorylation of MLKL’s pseudokinase domain by RIPK3 is the critical trigger in promoting MLKL disengagement from the necrosome, oligomerization, trafficking to the plasma membrane, and assembly into lytic hotspots where membrane permeabilization occurs^27, 32, 33, 34, 35, 36, 37^.

Because RIPK3-mediated phosphorylation of MLKL is a pivotal step in necroptotic signaling^29, 31^, several mouse models have been established to probe the functions of RIPK3^38, 39, 40, 41, 42, 43^. Deletion of RIPK3 does not compromise mouse viability and, in the absence of challenge, these mice are largely indistinguishable from wild-type counterparts^39, 40^. In comparison, many pathological processes are influenced by RIPK3 deficiency^20, 44^, although distinguishing whether these effects stem from RIPK3’s catalytic or non-enzymatic functions remains challenging. With the aim of deconvoluting this issue, three kinase-dead RIPK3 mouse strains have been generated: 1) The *Ripk3^K51A/K51A^* mouse strain harbors a mutation in the N-lobe of the kinase domain that eliminates canonical ATP-binding. Like *Ripk3^−/−^* mice, this strain is phenotypically normal under basal conditions^38^; 2) The *Ripk3*^D161N/D161N^ mouse strain carries a substitution in an activation loop residue that is required for Mg^2+^ binding. This strain has a fully penetrant embryonic lethal phenotype driven by errant Caspase-8-mediated apoptosis^41^; 3) The *Ripk3*^S165D/T166E^ mouse strain harbors two mutations in RIPK3’s activation loop that impair kinase activity and produces a phenotype similar to, albeit less severe than, *Ripk3*^D161N/D161N^ mice^42^. Why these kinase-inactivating mutations in RIPK3 yield different phenotypes remains unclear. It may indicate that the RIPK3^K51A^ mutation prevents both RIPK3’s necroptotic and apoptotic actions, whereas the RIPK3^D161N^ and RIPK3^S165D/T166E^ variants solely perturb necroptotic signaling. Understanding how RIPK3’s kinase domain regulates cell death remains a topic of great interest given that selective small inhibitors of RIPK3, while preventing necroptosis, also trigger on-target apoptosis^38^.

To further examine the role of RIPK3’s kinase domain in the necroptosis-to-apoptosis switch, we generated another knock-in strain harboring a conservative Asp-to-Asn substitution at residue 143 within the catalytic loop of RIPK3’s kinase domain, herein referred to as *Ripk3^D143N/D143N^* mice. In contrast to other kinase-dead RIPK3 variants, *Ripk3^D143N/D143N^* mice were born at a sub-Mendelian ratio due to the partial loss of homozygotes during embryogenesis. However, once born, *Ripk3^D143N/D143N^* mice were viable, fertile and developmentally normal into adulthood. The distinct phenotypes of the various kinase-inactivating mutations could be attributed to differences in their protein stability and binding repertoires. We observed that RIPK3^D143N^ was more stable than RIPK3^K51A^ and RIPK3^D161N^, and only RIPK3^D143N^ bound both RIPK1 and Caspase-8 in the presence of a caspase inhibitor. Overall, our data support the concept that the conformation of RIPK3’s kinase domain tightly regulates the higher-order assembly of a RIPK1- and Caspase-8-containing complex.

## RESULTS

### The kinase domain of RIPK3 controls apoptosis

RIPK3 catalytic activity is essential for necroptotic signaling because of its ability to phosphorylate and activate MLKL^29, 31^. RIPK3’s kinase activity relies on three active site residues: the ATP-positioning lysine within the β3 strand of the N-lobe (K51 in mouse RIPK3); the catalytic Asp of the catalytic loop HRD motif (D143); and the Mg^2+^ cofactor binding Asp of the DFG motif in the activation loop (D161) (**Fig. 1a**). To compare the functional impact of kinase-inactivating mutations at these sites, we stably transduced *Ripk3^−/−^* mouse dermal fibroblasts (MDFs) with lentiviral vectors encoding doxycycline-inducible wild-type RIPK3 (RIPK3^WT^), RIPK3^K51A^, RIPK3^D143N^ and RIPK3^D161N^. A concentration of 10 ng/mL doxycycline (dox) was chosen to reconstitute RIPK3 expression to endogenous levels (**Supp. Fig. 1A**). Unlike with RIPK3^WT^, reconstitution with RIPK3^K51A^, RIPK3^D143N^ and RIPK3^D161N^ mutants, did not restore sensitivity to the necroptotic stimulus, TNF/Smac mimetic/IDN-6556 (TSI; **Fig. 1Bi**), verifying that the mutants were catalytically inactive.

**Fig. 1.**
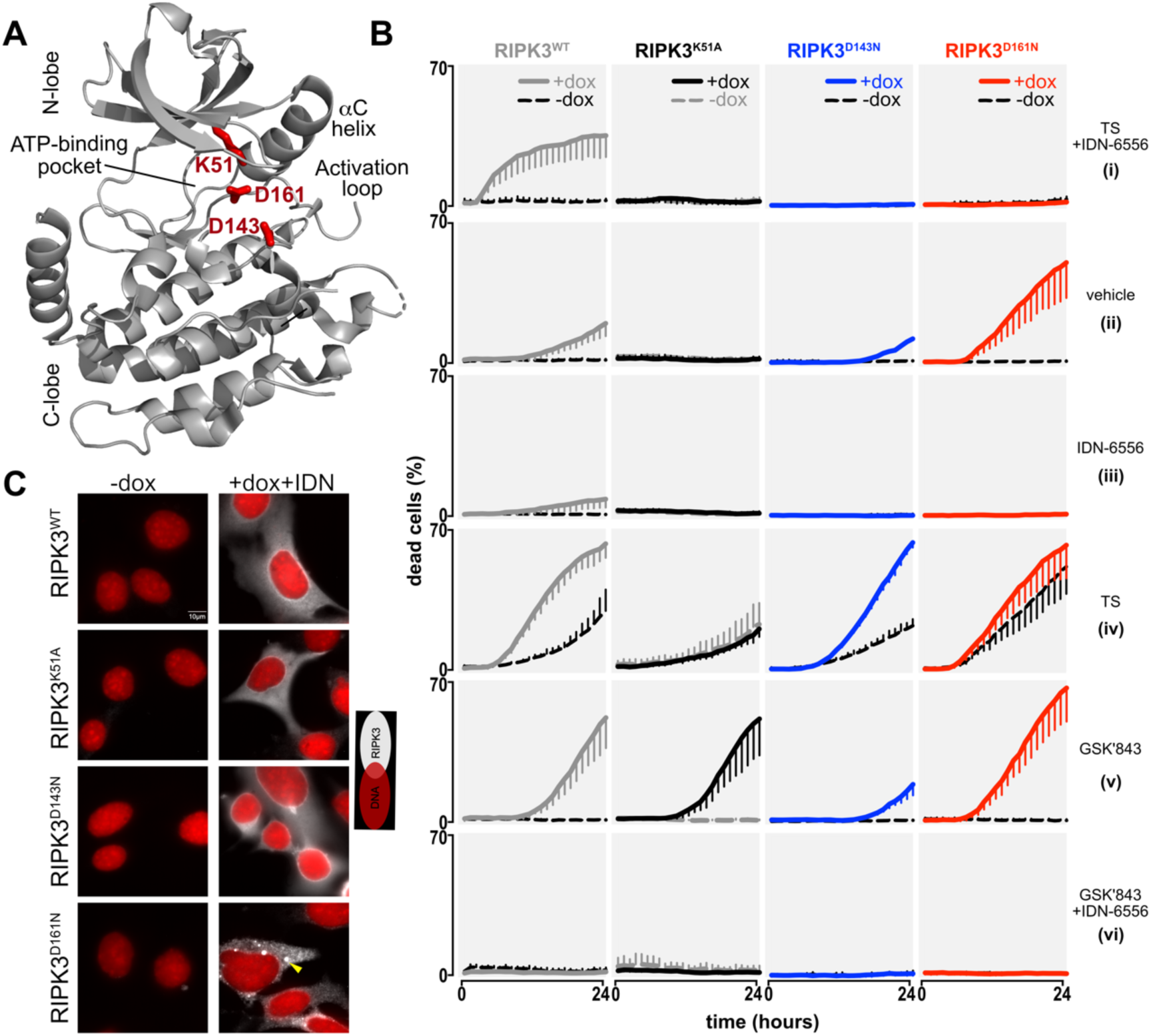
Cells expressing different kinase-dead RIPK3 mutants exhibit distinct sensitivities to apoptotic stimuli. **A** Richardson (Ribbons) diagram of the mouse RIPK3 kinase domain structure (PDB, 4M66)^62^ in which the three catalytic residues targeted for kinase dead mutations are shown as red sticks. **B** Death of *Ripk3^−/−^* MDFs without or with doxycycline to express RIPK3^WT^, RIPK3^K51A^, RIPK3^D143N^ or RIPK3^D161N^ in the presence of the stipulated stimuli. Treatments: (i) TNF, Smac mimetic and IDN-6556 (TSI); (ii) vehicle, DMSO; (iii) IDN-6556; (iv) TNF and Smac mimetic (TS); (v) GSK′843; (vi) GSK′843, IDN-6556. Mean±SEM for one experiment and representative of data from 2 independent experiments. Cell death calculated as the percentage of propidium iodide^+^ cells relative to the number of SPY700^+^ cells. **C** Immunofluorescence of *Ripk3^−/−^* MDF cells without or with doxycycline to express RIPK3^WT^, RIPK3^K51A^, RIPK3^D143N^ or RIPK3^D161N^. Data representative of two independent experiments. Red, Hoechst nuclear stain. White, RIPK3 immunosignal.

In the absence of an external stimulus, the reconstitution of *Ripk3^−/−^* MDF cells with RIPK3^WT^ or the RIPK3^K51A^, RIPK3^D143N^ and RIPK3^D161N^ mutants yielded different cell fates. RIPK3^K51A^ did not induce death, RIPK3^WT^ and RIPK3^D143N^ produced mild death, whereas RIPK3^D161N^ caused extensive death (**Fig. 1Bii**). These constitutive death responses were due to RIPK3-mediated apoptosis as they were prevented by the pan-Caspase inhibitor, IDN-6556 (**Fig. 1Biii**). Thus, consistent with prior reports^38, 41^, we found that RIPK3^D161N^ is a stimulus-independent trigger for apoptosis, whereas the other kinase-dead variants of RIPK3 were comparatively less toxic. We next studied the role of RIPK3 mutation on TNF-induced apoptosis. Interestingly, RIPK3^WT^ and RIPK3^D143N^, but not RIPK3^K51A^, increased the rate of apoptosis caused by TNF/Smac mimetic compared to cells in which exogene expression was not induced by dox (TS; **Fig. 1Biv**). Due to constitutive killing activity of RIPK3^D161N^, the impact of RIPK3^D161N^ on TNF-induced apoptosis could not be assessed. Thus, RIPK3’s role in TNF-induced apoptosis is blocked by the RIPK3^K51A^ mutation, but largely unaffected by the RIPK3^D143N^ mutation.

To further explore RIPK3’s apoptotic scaffolding function, we used GSK′843 – a selective RIPK3 inhibitor that targets the RIPK3 active site and induces Caspase-8-dependent apoptosis^38^. GSK′843 induced high levels of apoptosis in cells expressing RIPK3^WT^ and RIPK3^K51A^, but was well-tolerated by cells reconstituted with RIPK3^D143N^ (**Fig. 1Bv-vi**). Again, the influence of RIPK3^D161N^ on GSK′843-induced apoptosis could not be distinguished from its inherent toxicity. Thus, in contrast to TNF-induced apoptosis, RIPK3’s role in GSK′843-induced apoptosis is unaffected by the RIPK3^K51A^ mutation, but attenuated by the RIPK3^D143N^ mutation. Notably, the ability of RIPK3^D161N^ and GSK′843 to trigger apoptosis was reflected by their shared propensity to drive RIPK3 into large cytoplasmic puncta (**Fig. 1C** and **Supp. Fig. 1B**). Together, our data suggest that RIPK3’s kinase domain conformation finely tunes its apoptotic scaffolding function, with RIPK3^D161N^ constitutively inducing apoptosis, RIPK3^K51A^ obviating RIPK3’s involvement in TNF-induced apoptosis, and RIPK3^D143N^ attenuating GSK′843-induced apoptosis.

### Catalytically-dead RIPK3^D143N^ retains stability and RIPK1-Caspase-8-binding

We hypothesized that conformational differences among the three kinase-dead versions of RIPK3 underlies their variable stability and scaffolding properties. To address this possibility, we expressed each exogene in *Ripk3^−/−^* MDF cells, subjected each to a thermal gradient and examined RIPK3’s solubility by immunoblot. This assay was chosen on the basis that pro-apoptotic signaling by GSK′843 involves enhanced RIPK3 thermal stability (**Supp. Fig. 1C**). Interestingly, RIPK3^D143N^ was the only mutant that exhibited thermal stability comparable to wild-type RIPK3 (**Fig. 2A-B**), with the destabilization of RIPK3^K51A^ and RIPK3^D161N^ implying each is conformationally labile. These differences in the stability of RIPK3 had no bearing on the stability of endogenous MLKL (**Fig. 2A-B**), prompting us to examine which other interactions are modulated by RIPK3 mutation. Accordingly, we reconstituted *Ripk3^−/−^* MDF cells with RIPK3^WT^ or the kinase-dead mutants in the presence of IDN-6556 to prevent caspase-mediated cleavage of RIPK3^43, 45^, followed by a brief treatment with TSI to allow TNF-induced death complexes to form (**Fig. 2C**). Critically, the interactions of RIPK3^D143N^ resembled that of wild-type RIPK3, where it bound phospho-RIPK1 and Caspase-8 following necroptotic stimulation with TSI. On the other hand, RIPK3^K51A^ bound neither RIPK1 nor Caspase-8, which aligns with its inability to mediate TNF-driven cell death. While RIPK3^D161N^ was able to retain phospho-RIPK1 binding, it no longer precipitated with Caspase-8, which may underlie its constitutive killing activity. More broadly, these data imply that the stability and conformation of RIPK3’s kinase domain, in addition to its catalytic activity, is a key determinant of apoptotic and necroptotic signaling.

**Fig. 2.**
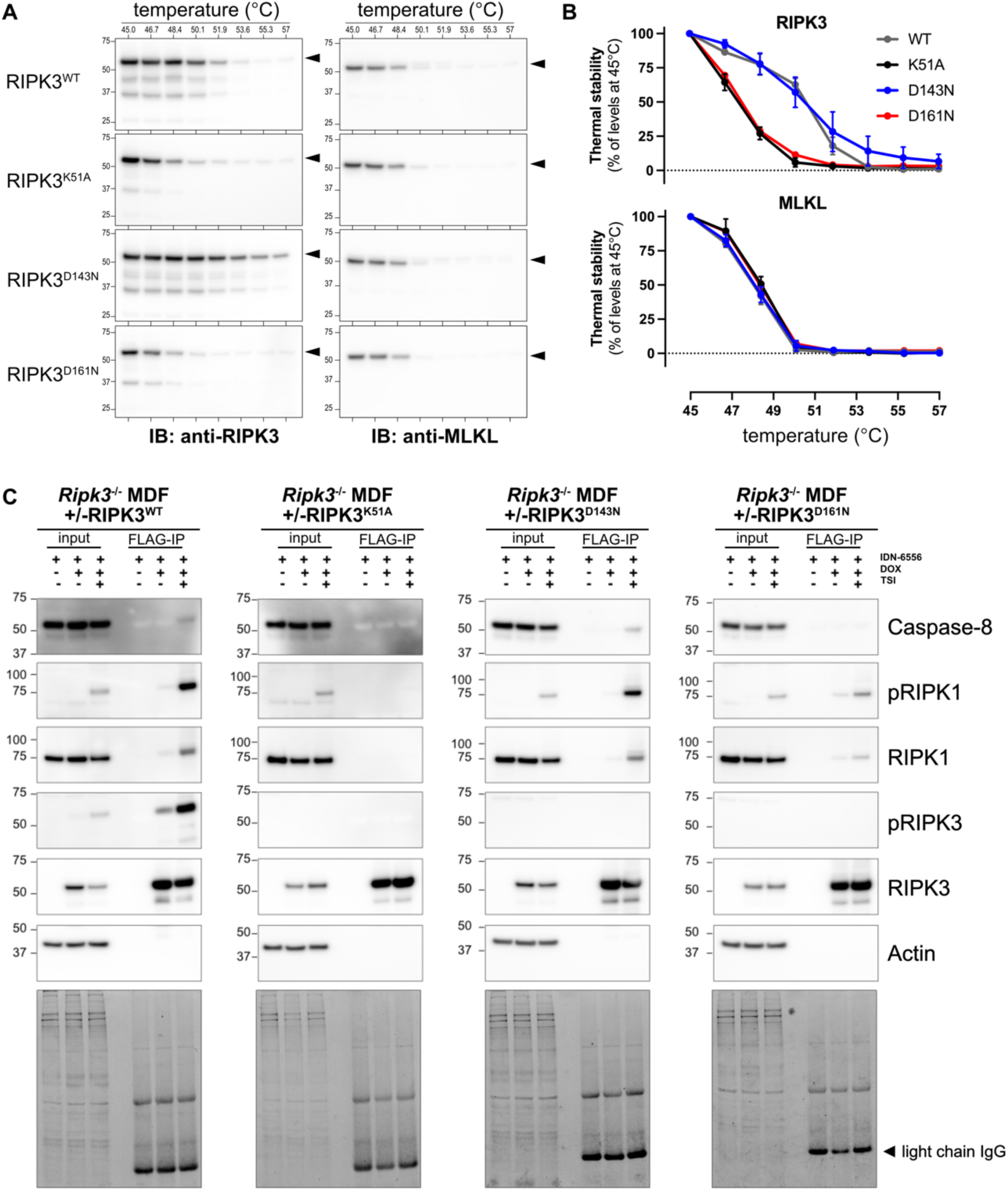
RIPK3^D143N^ retains stability and ability to bind to Caspase-8 and RIPK1. **A-B** Immunoblots (Panel A) and quantitation (Panel B) of cellular thermal stability assay of RIPK3^WT^, RIPK3^K51A^, RIPK3^D143N^ and RIPK3^D161N^. Representative of n=4 for RIPK3^WT^, RIPK3^K51A^, RIPK3^D143N^, and n=1 for RIPK3^D161N^. Mean±SEM. **C** Immunoblot of input and co-immunoprecipitations (FLAG-IP) of reconstituted RIPK3^WT^, RIPK3^K51A^, RIPK3^D143N^ and RIPK3^D161N^ in *Ripk3^−/−^* MDFs in the presence of IDN-6556 ± overnight doxycycline (DOX), then 2 hours of necroptotic stimuli (TSI). Data representative of n=2.

### Adult Ripk3^D143N/D143N^ mice are viable, fertile and ostensibly normal

In contrast to RIPK3^K51A^ and RIPK3^D161N^ mutants, RIPK3^D143N^ retains RIPK3^WT^-like stability and cellular interactions, making it an ideal kinase-dead mutant to explore RIPK3’s non-necroptotic actions *in vivo*. Accordingly, we generated a mouse harboring a RIPK3^D143N^-encoding mutation within exon 3 of the mouse *Ripk3* gene using CRISPR/Cas9 editing (**Fig. 3A**). *Ripk3^D143N/D143N^* mice were viable, fertile homozygous knock-in animals (**Fig. 3B**) that developed normally into adulthood (**Supp. Fig. 2A-G**). Intriguingly, while heterozygous matings produced a Mendelian distribution of genotypes at embryonic day 10 (E10.5), *Ripk3^D143N/D143N^* mice were slightly but significantly under-represented at weaning (**Fig. 3C**; χ^2^ test for goodness of fit showed p=0.88 at E10.5 and p=0.02 at weaning). This observation suggests that *Ripk3^D143N/D143N^* mice are partially sensitized to the same peripartum checkpoint that causes lethality in *Ripk3^D161N/D161N^* mice and *Ripk3*^S165D/T166E^ mice^41, 42^. In unchallenged adult *Ripk3^D143N/D143N^* mice, RIPK3^D143N^ was expressed and distributed comparably to wild-type RIPK3 in all examined tissues (**Fig. 3D** and **Supp. Fig.** 3), consistent with the stability of exogenous RIPK3^D143N^ observed in our cellular studies. Importantly, MDF cells and bone marrow-derived macrophages from *Ripk3^D143N/D143N^* mice confirmed that the endogenously-expressed RIPK3^D143N^ was catalytically inactive because these cells were completely protected from TNF-induced necroptotic death yet could still undergo TNF-induced apoptosis (**Fig. 3E-F**). Collectively, these data show that endogenous RIPK3^D143N^, despite conferring a transient and partial toxicity *in utero*, is stable and well-tolerated in mouse cells and tissues.

**Fig. 3.**
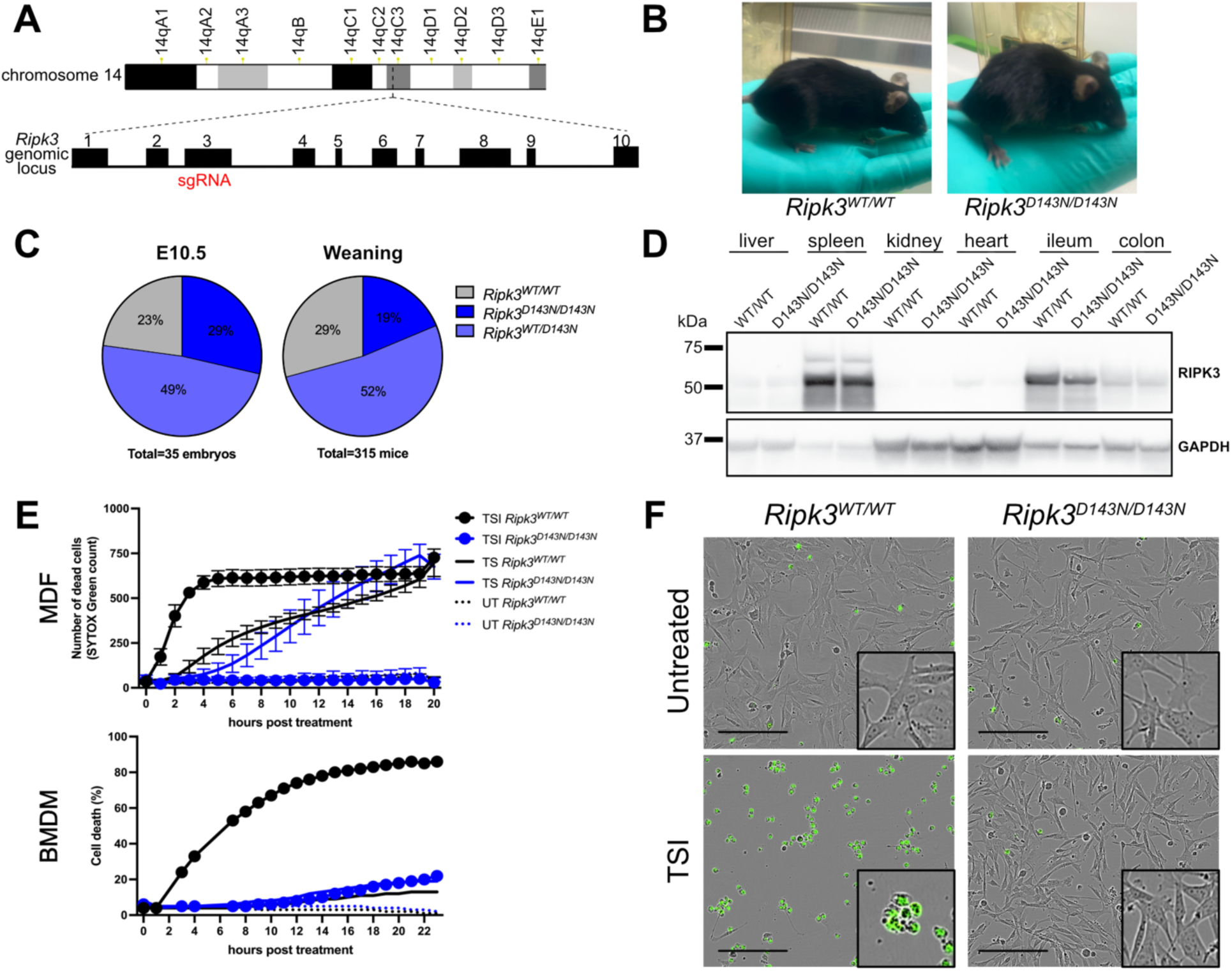
*Ripk3^D143N/D143N^* mice resemble *Ripk3^WT/WT^* littermates in the absence of challenge. **A** CRISPR gene editing strategy for the generation of *Ripk3^D143N/D143N^* mice. **B** *Ripk3^WT/WT^* and *Ripk3^D143N/D143N^* littermates at 4 months-of-age. **C** Genotype distribution at E10.5 or at weaning from heterozygous *Ripk3^D143N/WT^* matings. Statistical test to compare observed versus expected genotype ratios: χ^2^ test for goodness of fit showed p=0.88 at E10.5 and p= 0.02 at weaning. **D** Immunoblot of organ homogenates of littermates, n=2 per genotype. **E** Quantitation of the death of mouse dermal fibroblasts (MDF) and bone marrow derived macrophages (BMDM) from *Ripk3^WT/WT^* and *Ripk3^D143N/D143N^* mice. Cells were untreated (UT) or stimulated with TNF+Smac mimetic (TS) or TS+IDN-6556 (TSI). Cell death was measured by the uptake of SYTOX Green. BMDM death was calculated as the percentage of SYTOX Green^+^ cells relative to the number of SPY620^+^ cells. MDF data representative of two independent experiments across three independent cell lines. BMDM data representative of one experiment. **F** Micrographs of *Ripk3^WT/WT^* and *Ripk3^D143N/D143N^* MDFs after 19 hours of treatment. Scale bars represents 200 µm.

### RIPK3^D143N^ protects mice from MLKL-dependent intestinal inflammation

We next sought to determine whether endogenous RIPK3^D143N^ protects from *in vivo* necroptosis. To this end, we established a colony of *Villin*^1000^–Cre *Casp8^fl/fl^* mice (herein referred to as *Casp8^IEC^* mice) where the Villin promoter drives Cre expression to delete Caspase-8 from intestinal epithelial cells, which in turn triggers RIPK1-RIPK3-MLKL-dependent Paneth cell loss and intestinal inflammation^20, 21, 46^. We confirmed that Villin^1000^–driven Cre activity was on-target (**Fig. S4A**; mild off-target Cre activity was found in the liver) and induced selective and complete deletion of Caspase-8 from the intestinal epithelia in *Casp8^IEC^* mice (**Supp. Fig. 4B**). Consistent with prior reports^20, 21, 46^, *Casp8^IEC^* mice developed endoscopically-apparent colitis (**Fig. 4A-B**), with a significant proportion of the Lyzozyme^+^ and MPTX2^+^ Paneth cells lost by 6 months-of-age (**Fig. 4C-D** and **Supp. Fig. 5A-B**). Beyond these manifestations, *Casp8^IEC^* mice in our facility exhibited a mild phenotype without impacts on survival (**Supp. Fig. 5C**), weight (**Supp. Fig. 5D**), or Goblet cell numbers (**Fig. 4E-F**). Importantly, crossing *Casp8*^IEC^ mice onto the *Ripk3^D143N/D143N^* background completely prevented Paneth cell loss (**Fig. 4C-D**). Similarly, *Casp8^IEC^Mlkl^IEC^* mice with conditional deletion of both caspase-8 and MLKL were fully protected from Paneth cell loss (**Fig. 4C-D**). Altogether, these data show, for the first time, that cell-intrinsic necroptosis is solely responsible for Paneth cell loss in the *Casp8^IEC^* model. Why Paneth cells, which express low levels of RIPK3 under basal conditions^14^, are the only intestinal epithelial population that succumbs to Caspase-8-deficiency remains unclear. Interestingly, intestinal epithelial RIPK3 expression was increased in *Casp8^IEC^* mice, but remained at basal levels in *Casp8^IEC^Ripk3^D143N/D143N^* and *Casp8^IEC^Mlkl^IEC^* mice (**Fig. 4G**). Because intestinal RIPK3 levels increase with inflammation^14^, this finding indicates that Paneth cell necroptosis is the precipitating event that promotes downstream intestinal inflammation. More broadly, our study shows that the *Ripk3^D143N^* mouse model is an elegant way of preventing necroptosis *in vivo*, and provides definitive support for the crucial interplay of Caspase-8 and the necroptotic machinery in provoking gut inflammation.

**Fig. 4.**
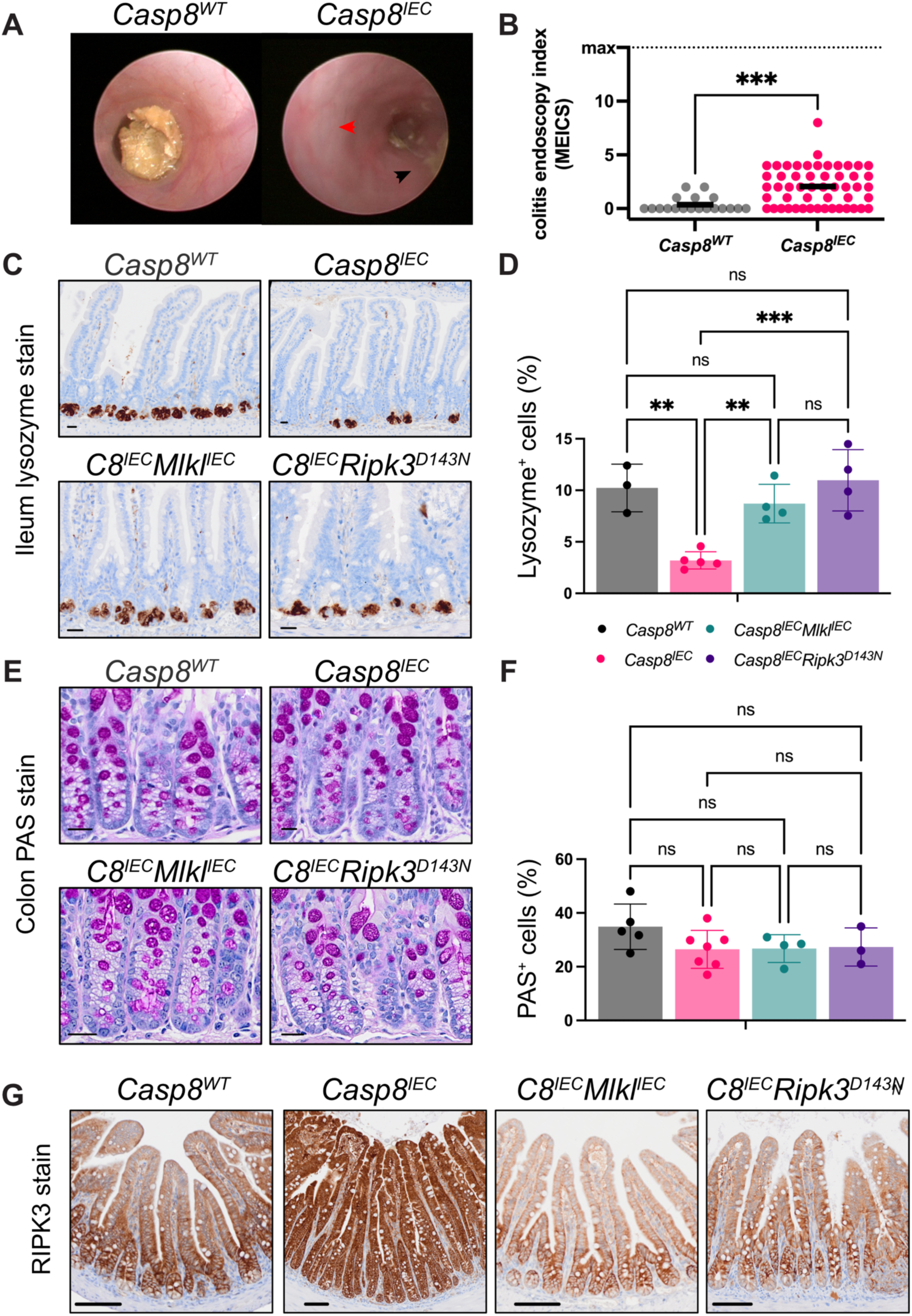
Caspase-8-deficient Paneth cells survive in *Ripk3*^D143N/D143N^ mice. **A** Endoscopy images of a healthy wild-type (*Casp8*^WT^) mouse and a *Casp8^IEC^* mouse with colitis. Red arrow points to the loss of vasculature and black arrow points to a loose stool. **B** Murine endoscopic index of colitis severity (MEICS) scores for *Casp8*^WT^ and *Casp8^IEC^* mice. Each dot represents the score from a mouse during their monthly endoscopy session between 6-24 weeks-of-age. Line indicates mean. ***p<0.001 from unpaired non-parametric t-test with Mann-Whitney correction. *Casp8^WT^* n=13 mice. *Casp8^IEC^* n=12 mice. **C** Periodic Acid-Schiff (PAS; magenta) staining. Scale bars are 20 µm. Images representative of *Casp8^WT^ n*=5, *Casp8^IEC^ n*=6, *Casp8^IEC^Mlkl^IEC^ n*=4, and *Casp8^IEC^Ripk3^D143N/D143N^ n*=3 mice. **D** PAS^+^ cells as a percentage of total cells. Same cohort as in Panel E. An average of 70,000 cells were measured per mouse with each dot representing the mean value from one mouse. Bars indicate group mean ±SD. n.s. indicates p>0.05 by ordinary one-way ANOVA with Tukey’s correction. **E** Immunohistochemistry (brown) signals for lysozyme. Scale bars are 20 µm. Images representative of *Casp8^WT^* n=3, *Casp8^IEC^* n=4, *Casp8^IEC^Mlkl^IEC^* n=3, and *Casp8^IEC^Ripk3^D143N/D143N^* n=4 mice. **F** Lysozyme^+^ cells as a percentage of total cells. Same cohort as in Panel C. An average of 35,000 cells were measured per mouse with each dot representing the mean value from one mouse. Bars indicate group mean ±SD. *p<0.05 and **p<0.01 by ordinary one-way ANOVA with Tukey’s correction. **G** Immunohistochemistry signals for RIPK3 in the ileum of *Casp8^WT^*, *Casp8^IEC^*, *Casp8^IEC^Mlkl^IEC^*, and *Casp8^IEC^Ripk3^D143N/D143N^* mice; representative of *n*=3, 5, 3, and 4 mice. Scale bars are 100 µm.

## DISCUSSION

Until recently, observations that RIPK3 inhibition can promote apoptosis have raised concerns about RIPK3’s viability as a therapeutic target^38, 41^. However, subsequent findings indicate that RIPK3 inhibitors are well-tolerated in animals^47, 48, 49^, suggesting RIPK3 kinase domain conformation, rather than loss of catalytic activity, is likely to underpin the previously reported Caspase-8-driven toxicity. Here, we tested this idea by generating a knock-in mouse strain harboring the RIPK3^D143N^ kinase-dead mutant. The resulting *Ripk3^D143N^* homozygote mice were viable, fertile, with no detectable pathology or hematopoietic defects in the absence of challenge, unlike the embryonic lethality found in *Ripk3^D161N^* strain^41^. Collectively, our data support the idea that RIPK3^D143N^ is akin to RIPK3^WT^ in interactions and stability, leading us to propose that *Ripk3^D143N/D143N^* mice should be adopted as the preferred model for studying the non-catalytic roles of RIPK3 *in vivo*. We demonstrate the utility of the *Ripk3^D143N/D143N^* strain as a model for studying necroptosis *in vivo* by showing protection from the ileitis caused by intestinal epithelial cell knockout of *Casp8*. The protection conferred by the RIPK3^D143N^ mutation was phenocopied by deletion of *Mlkl* specifically within intestinal epithelial cells, indicating that blockade of necroptosis within intestinal epithelial cells, rather than infiltrating cells, confers protection.

While adult *Ripk3^D143N/D143N^* mice grossly resembled wild-type counterparts, we observed a partially-penetrant embryonic lethality manifesting in sub-Mendelian ratio of homozygotes at weaning. Because embryonic lethality was previously observed for the *Ripk3^D161N^* strain^41^ and *Ripk3^S165D/T166E^* strain^42^, but not *Ripk3^K51A^* mice^38^, it was known that loss of RIPK3 catalytic activity itself is not deleterious to development. Here, our findings with the *Ripk3^D143N^* knock-in mouse support this assertion, and instead indicate that the conformation of the RIPK3 kinase domain is a crucial determinant of cell, and organism viability. Much emphasis has been placed on the role of the RIPK1 and RIPK3 RHIMs in assembling the necrosome. Whilst it has been long-established that RIPK1 autophosphorylation of S166 within its kinase domain is crucial to propagation of necroptotic signals^18, 50, 51^, our data support a model in which the conformation of the RIPK3 kinase domain allosterically influences RIPK3’s interaction with RIPK1, which in turn influences downstream cell death signaling (**Fig. 5**). How this allosteric regulation extends to Caspase-8 sequestration and activation, and impacts the stoichiometry and connectivity of RIPK3-containing complexes, remains a major focus for future investigation.

**Fig. 5.**
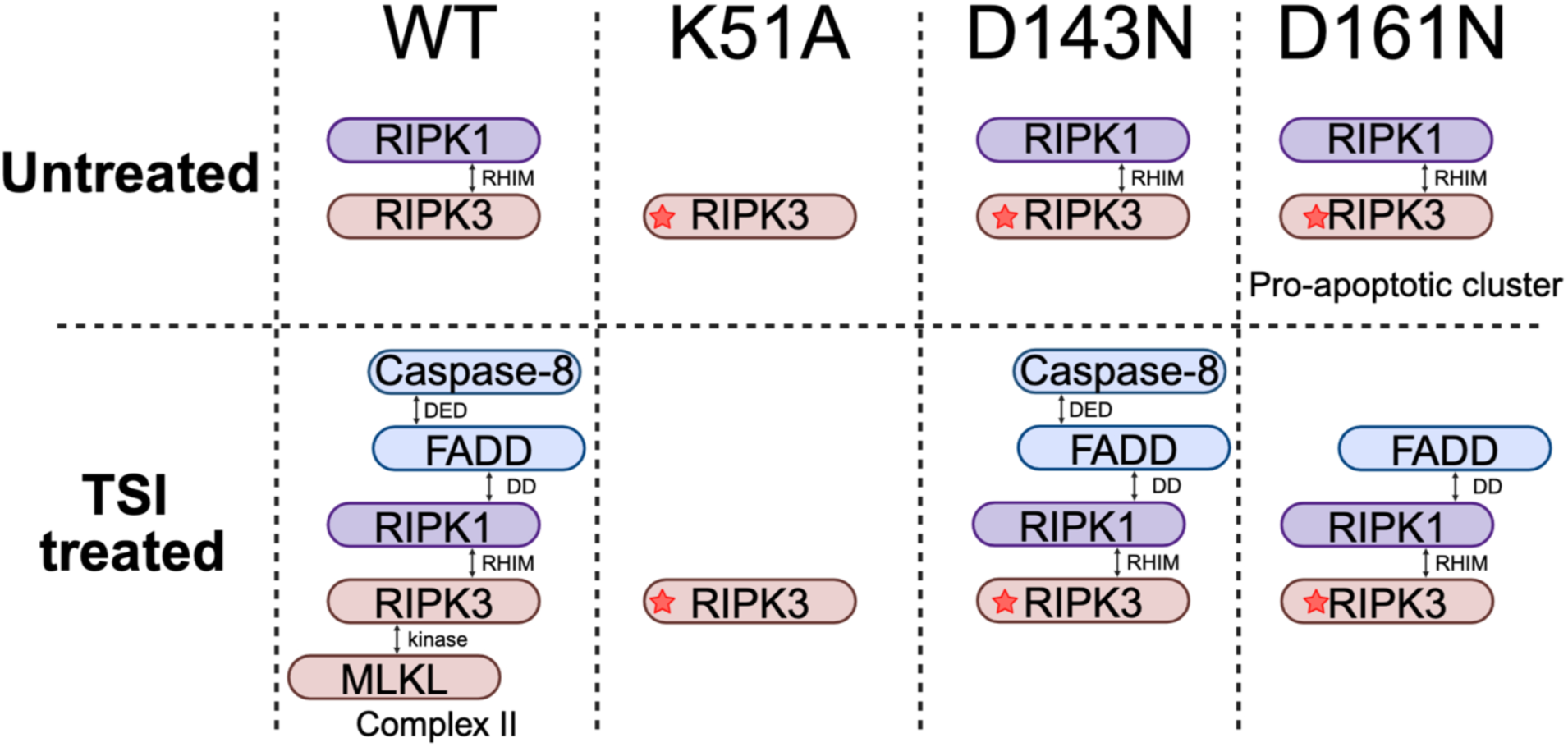
Working model for the cellular interactions of wild-type RIPK3 and its kinase-dead mutants. Star indicates mutant form of RIPK3. RIP homotypic interaction motif (RHIM), Death Effector Domain (DED), Death Domain (DD).

## METHODS

### Mice

Mice were housed at the Walter and Eliza Hall Institute of Medical Research (WEHI), Australia. Animal experiments were approved by the WEHI Animal Ethics Committee (2022.085) in accordance with the Prevention of Cruelty to Animals Act (1986) and the Australian National Health and Medical Research Council Code of Practice for the Care and Use of Animals for Scientific Purposes (1997). Mice were housed in a temperature and humidity controlled specific pathogen free facility with a 12 hour:12 hour day night cycle. Villin.cre1000 mice were purchased from Jackson Laboratories (Jackson Laboratories, Maine, United States; Strain #021504). The floxed *Casp*8 mouse strain^52^, the floxed *Mlkl* mouse strain^31^ and the *Rosa26* Ai9 Cre reporter mouse strain^53^ have been reported previously. All mice in this study were maintained on a C57BL/6 J background. Co-housed littermate mice, rather than randomized cohorts, were preferentially used for the *Casp8^IEC^* model. Unless stipulated, mice in the *Casp8^IEC^* model were harvested at 5-6 months of age.

### Generation and genotyping of the Ripk3^D143N/D143N^ mouse strain

*Ripk3^D143N/D143N^* mice on C57BL/6J background were generated with CRISPR/Cas9 by the Melbourne Advanced Genome Editing Centre (MAGEC) laboratory. The *Ripk3* gene on mouse chromosome 14 was targeted to induce the D143N mutation. One sgRNA of the sequence TGTTAGAGGGCTTGAGGTCC was used to create double stranded breaks within the *Ripk3* locus to stimulate homologous recombination. An oligonucleotide donor with the sequence CCTGCTGCAGGAAGTGGTGCTGGGGATGTGCTACCTACACAGCTTGAACCCTCCGCTCCTGCACCGGAATCTCAAGCCCTCTAACATTCTGCTGGATCCAGAGCTCCACGCCAAGGTTAGTCCATCTACA was used to introduce the D143N mutation. To detect the integration of the donor, mice were genotyped using the forward primer GGGACAGGTACTCAACATGGTC and the reverse primer TTTCGGCGTGTAGGAAGAAG, with sequencing overhangs attached. The PCR conditions for genotyping were: 95°C 2 minutes, (95°C 30 seconds, 60°C 30 seconds, 72°C 30 seconds) x 35, 72°C 5 minutes. F0 mice were backcrossed with wildtype C57BL/6J mice for at least 2 generations to breed out any potential off-target mutations.

### Immunoblotting

Cells/tissues were lysed in ice-cold RIPA buffer (10 mM Tris-HCl pH 8.0, 1 mM EGTA, 2 mM MgCl_2_, 0.5% v/v Triton X-100, 0.1% w/v sodium deoxycholate, 0.5% w/v sodium dodecyl sulfate (SDS), and 90 mM NaCl) supplemented with 1× Protease and Phosphatase Inhibitor Cocktail (Cell Signaling Technology Cat#5872) and 100 U/mL Benzonase (Sigma-Aldrich Cat#E1014) to a concentration of 50 mg/mL (w/v) with a stainless-steel bead using a Qiagen TissueLyser II (1 minute at 30 Hz). Homogenates were boiled for 10 minutes in 1 × SDS sample buffer (126 mM Tris-HCl, pH 8, 20% v/v glycerol, 4% w/v SDS, 0.02% w/v bromophenol blue, 5% v/v 2-mercaptoethanol) and ran on 1.5 mm NuPAGE 4–12% Bis-Tris gels (ThermoFisher Scientific NP0335BOX) in MES Running Buffer (ThermoFisher Scientific NP000202) at 150 V for 60 minutes. Gels were transferred to a polyvinylidene difluoride membrane (Merck Cat# IPVH00010) at 100 V for 60 minutes and then blocked in 5% w/v skim milk powder in TBS-T (50 mM TrisHCl pH 7.4, 0.15 M NaCl, 0.1 v/v Tween-20). After transfer, gels were stained with SimplyBlue SafeStain (ThermoFisher Scientific LC6065) and membranes were probed with the following antibodies: RIPK3 (clone 8G7; ref.^8^; produced in-house; available from Millipore as MABC1595), RIPK3 (clone 1H12; ref.^54^; produced in-house), MLKL (clone 3H1; ref.^31^; produced in-house; available from Millipore as MABC604), Caspase-8 (clone F5K9P; Cell Signaling Technology), RIPK1^pS166^ (Cat#31122; Cell Signaling Technology), RIPK1 (clone D94C12; Cell Signaling Technology), RIPK3^pS231pS232^ (clone E7S1R; Cell Signaling Technology), RIPK3 (clone D4G2A; Cell Signaling Technology), horseradish peroxidase (HRP)-conjugated Actin (Santa Cruz Biotechnology, Cat# sc-47778), and GAPDH (Millipore MAB374). Primary antibodies produced in-house were used at 1 μg/mL for immunoblot, and all other primary antibodies were used at a 1:1000-2000 dilution in Tris-Balanced Salt Solution containing 0.1% Tween 20 (TBS+T) supplemented with 5% w/v cow’s skim milk powder and 0.01% w/v sodium azide. Primary antibody probes were performed overnight at 4 °C on a rocker before washing twice in TBS+T, then membranes were probed with a 1:10000 dilution of HRP-conjugated secondary antibody: goat anti-rat immunoglobulin (Ig; Southern BioTech Cat#3010-05), goat anti-rabbit Ig (Southern BioTech Cat#4010-05), and goat anti-mouse Ig (Southern BioTech Cat#1010-05). Membranes were washed four times in TBS+T and signals revealed by enhanced chemiluminescence (Merck Cat#WBLUF0100) on a ChemiDoc Touch Imaging System (Bio-Rad). Between probing with primary antibodies from the same species, membranes were incubated in stripping buffer (200 mM glycine pH 2.9, 1% w/v SDS, 0.5 mM TCEP) for 30 minutes at room temperature and then re-blocked. Uncropped immunoblots are included as a Supplemental Data file.

### Immunohistochemistry (IHC)

Mice were euthanized via gradual carbon dioxide asphyxiation, with bipedal and orbital reflexes checked to ensure death. Tissues were harvested and immediately placed in 10% Neutral-Buffered Formalin at a 1:10 tissue:formalin ratio at room temperature. Formalin fixed tissues were paraffin embedded using the standard 8-hour auto-processing protocol of Tissue-Tek VIP® 6 AI Tissue Processor (Sakura Finetek USA) within 1-2 days of harvest. Four-micron-thick sections of paraffin-embedded tissues were cut onto adhesive slides (Menzel Gläser Superfrost PLUS). IHC and image acquisition of sections stained for Caspase-8, RIPK1, RIPK3 and cleaved Caspase-3 followed protocols described previously^55^; staining for lysozyme (Abcam, Cat#108508) and for MPTX2 (Abcam, Cat#EPR20920-19; marker of Paneth cells^56^) followed the protocols detailed in Supplementary Table 1.

### Cellular thermal stability assay

*Ripk3^−/−^* mouse dermal fibroblasts with doxycycline-inducible expression constructs for RIPK3^WT^, RIPK3^K51A^, RIPK3^D143N^ or RIPK3^D161N^ were seeded to ∼50% confluency. After 2 hours, cells were treated with 10 ng/mL doxycycline and 5 μM IDN-6556. The next day, cells were trypsinized, pelleted (700 ×*g*, 5 minutes, room temperature), then resuspended in dPBS supplemented with cOmplete EDTA-free protease inhibitor (Merck, Cat#04693132001) and PhosSTOP phosphatase inhibitor (Merck, Cat#4906837001). 100 μL of 2×10^6^ cells/mL were aliquoted into PCR tubes and incubated across a gradient of 8 temperatures (45-57°C) for 3 minutes, then at 21°C for 3 minutes, then at 12°C for 1 minute. Samples were immediately frozen at −80°C, thawed on ice, pelleted (20000 ×*g*, 20 minutes, 4°C) and 35 μL of supernatant analyzed via immunoblot. Immunoblot signals were quantified with Image Lab v6.1 (Bio-Rad) using the raw full-resolution .scn Chemidoc files.

### Immunoprecipitation

*Ripk3^−/−^* mouse dermal fibroblasts with doxycycline-inducible expression constructs for RIPK3^WT^, RIPK3^K51A^, RIPK3^D143N^ or RIPK3^D161N^ were seeded into 6-well plates at 2×10^5^ cells/well. 2 hours later, cells were treated with 5 μM IDN-6556 and ± 10 ng/mL doxycycline. The next day, cells were treated with 100 ng/mL tumor necrosis factor, 500 nM Smac mimetic Compound A and 5 µM IDN-6556. 90 minutes later, cells were washing with PBS, then lysed with 1 mL/well of lysis buffer (50 mM Tris-HCl, pH 7.4, 1 % v/v Triton X-100, 150 mM NaCl, cOmplete EDTA-free protease inhibitor (Merck, Cat#04693132001), PhosSTOP phosphatase inhibitor (Merck, Cat#4906837001) and 100 U/mL Benzonase (Sigma #E1014). Lysates were freeze-thawed and a 70 µL aliquot kept aside as the “input”. The remainder of each lysate was supplemented with 20 µL/sample of anti-HA magnetic beads (ThermoFisher Scientific Cat#88837), samples rotated (30 minutes, 4°C), beads pelleted and discarded, 120 µL/tube of anti-Flag magnetic beads (Merck, Cat#M8823) was added to the supernatant, samples rotated (90 minutes, 4°C), beads pelleted and washed three times with 0.5 mL of lysis buffer, beads pelleted, resuspended in 75 µL of 2xSDS loading buffer, incubated (10 minutes, 100°C), beads pelleted and the supernatant analyzed via immunoblot.

### Assessments of colitis severity

Colonoscopies on mice were performed every four weeks from 6-24 weeks of age using established method^57, 58^. Briefly, mice were anaesthetized via the gradual increase of isoflurane (0.5-5% v/v), pedal reflex was checked to ensure deep anesthesia, the endoscope probe inserted into the rectum (Coloview miniendoscopic system including Endovision Tricam (Karl Storz Cat#20212001-020); Xenon 175 light source with anti-fog pump (Karl Storz Cat#20134001); HOPKINS straight Forward Telescope (Karl Storz Cat#64301A); Endoscopic Sheath (total diameter 3 mm; Kalr Stroz Cat#61029C); Fibre Optic Light Cable (Kalr Stroz Cat#69495ND)) and then a video recorded with computer and media player software (Apple Inc., iMovie). Colonoscopy videos were scored using the murine endoscopic index of colitis severity (MEICS) rubric^57^ in a blinded fashion.

### Mouse dermal fibroblasts and bone marrow derived macrophages culturing

Mouse dermal fibroblasts (MDFs) were prepared from the skin of mouse tails then immortalized via stable lentiviral transduction with SV40 large T antigen as described previously^31^ and maintained in DMEM + 8% fetal calf serum. Bone marrow-derived macrophages (BMDMs) were generated from the marrow harvested of mice femurs. Harvested cells were maintained in DMEM supplemented with 20% L929 conditioned media and 8% fetal calf serum for at least 7 days before experimentation. All cells were maintained at 37℃ under humidified conditions and 10% v/v carbon dioxide.

### Induction and quantification of cell death

MDFs and BMDMs were seeded at 20,000 cells/well and 120,000 cells/well in 48-well plates, respectively. The next day, cells were treated with stimuli in DMEM supplemented with 1:20000 SYTOX Green (Invitrogen, Cat#S7020) ± 1:1000 SPY620-DNA (Spirochrome, Cat#SC401) or in DMEM supplemented with 1:2000 propidium iodide (ThermoFisher Scientific; Cat#P3566) ± 1:2000 SPY700-DNA (Spirochrome, Cat#SC601). The stimuli used were: 10 ng/mL doxycycline, 5 μM IDN-6556, 10 μM GSK′843, 100 ng/mL tumour necrosis factor, 500 nM Smac mimetic compound A. Directly after stimulation, cells were moved into an IncuCyte SX5 System (Essen Bioscience) and imaged using the 10x objective over time using the default bright-field, green, red and far-red channel settings. SYTOX Green^+^ or propidium iodide^+^ dead cells were quantified using IncuCyte SX5 v2022B software (Essen Bioscience).

### Immunofluorescence and image acquisition

*Ripk3^−/−^* mouse dermal fibroblasts with doxycycline-inducible expression constructs for RIPK3^WT^, RIPK3^K51A^, RIPK3^D143N^ or RIPK3^D161N^ were seeded at 15,000 cells/well into 8-well chamber slides (Ibidi Cat#80826). 2 hours later cells were treated with 10 ng/mL doxycycline and 5 μm IDN-6556, then 9 hours later cells were treated ± 10 μM GSK′843. 90 minutes later, slide was ice-chilled for 3 minutes, then washed in ice-cold dPBS, then fixed for 15 minutes in ice-cold methanol. Cells were washed once in ice-cold dPBS, then blocked in ice-cold Tris-balanced salt solution with 0.05% v/v Triton-×100 (TBS+T) supplemented with 10% v/v donkey serum (Sigma #D9663) for >1 hour. Cells were incubated in 2 μg/mL anti-RIPK3 antibody (clone 1H12; ref.^54^; produced in-house) overnight at 4 °C in TBS+T with 10% v/v donkey serum. Cells were washed twice in TBS+T then incubated in 1:10000 AlexaFluor488-conjugated donkey anti-rat IgG supplemented with 0.1 μg/mL Hoechst 33342 (ThermoFisher Scientific Cat#H3570) for 3 hours at room temperature with gentle rocking. Cells were washed four times in ice-cold TBS+T then stored at 4 °C until being imaged on an Inverted Axio Observer.Z1 microscope (Zeiss) with the following specifications: Plan-Apochromat 100x/1.4 Oil Ph3 M27 lens, HXP 120 V excitation source, AlexaFluor488 imaged with a λ_Excitation_ = 450–490 nm; λ_beamsplitter_ = 495 nm; λ_Emission_ = 500– 550 nm, Hoechst 33342 imaged with a λ_Excitation_ = 359–371 nm; λ_beamsplitter_ = 395 nm; λ_Emission_ = 397-∞ nm, a sCMOS PCO.edge 4.2 camera, ZEN blue v3.10.103 capture software and ImageJ 1.54p post-acquisition processing software^59^. Typically, for each independent experiment, 5–10 randomly selected fields were captured per treatment group, where only the Hoechst 33342 signal was visualised prior to multi-channel acquisition. To ensure consistent signal intensities across independent experiments, the same excitation, emission and camera settings were used throughout this study.

### Histopathology of mice

Comprehensive full body necropsy, Haematoxylin and eosin (H&E) images, blood analysis reports of wildtype *n*=3 and *Ripk3^D143N^ n*=3 littermates between 7-10 months of age were prepared by veterinary and medical pathologists from Phenomics Australia, Melbourne Victoria.

### Hematological analysis

Submandibular bloods of mice from 2-13 months of age were collected into EDTA-coated tubes (Sarstedt Microvette® 500 EDTA K3E, 500 µL, Order number: 20.1341). Blood samples were diluted by a factor of 1-13-fold in dPBS and blood cells quantified with ADVIA 2120 hematological analyzer on the same day as blood collection.

### Periodic acid–Schiff stain

The manual staining protocol for Periodic acid–Schiff stain (PAS, Sigma-Aldrich, Cat#1090330500) is detailed in Supplementary Table 1.

### Acquisition of (immuno)histochemistry images

Stained slides were scanned on: Olympus VS200 (objective: 20x, numerical aperture 0.8, media dry; software: Olympus VS200 ASW 3.41). Representative full-resolution 8-bit RGB micrographs of tissues were imported into ImageJ 1.53t^59^. Capture settings and post-acquisition image transformations were held constant between any micrographs that were being compared.

### Assessment of Villin.Cre specificity

Fresh frozen tissue sections from *ROSA26 Ai9* Villin.Cre1000 mice and wild-type control mice were mounted in DAPI-containing media and imaged on a Vectra Polaris Imaging System (Akoya Biosciences), as described previously^60^. Acquisition software: Vectra Polaris v.1.0. Resolution: 0.5 μm/pixel (20x objective). LED light source. Default filters for DAPI MSI (4 seconds exposure) and Texas Red (150 seconds exposure) were used.

### Quantification of cells by IHC

Full resolution scans of IHC stained tissue sections were opened in QuPATH software v.0.5.1. In brief, a selection tool is used to highlight an area of an average of 50% of Swiss rolls. IHC signal was unmixed from Hematoxylin using the “Color Deconvolution 2” plugin^61^. IHC-stain positive cells were segmented and quantified. Percentage of PAS^+^ cells as total cells were calculated with two scripts, “PAS^+^ cell detection” and “Total cell count”. Percentage of Lysozyme^+^ or MPTX2^+^ cells were calculated with “Lysozyme^+^ cell detection and total cell count”. “Threshold” for each script was adjusted with each staining batch due to natural variations, and tissue sections from mouse models of IBD were accompanied by wildtype littermate control on the same slide. The scripts are provided in Supplementary Table 2.

### Statistical analysis

The number of independent experiments for each dataset is stipulated in the respective figure legend. The employed statistical tests are stipulated in the respective figure legend and performed using Prism v.10 (GraphPad).

## DATA AVAILABILITY

The uncropped blots from this study are included as Supplementary Data. Other raw data from this study are available from the corresponding authors upon request.

## MATERIALS AVAILABILITY

Any materials from this study that are not commercially available will be provided under Materials Transfer Agreement upon request to the corresponding authors.

## ACKNOWLEDGEMENTS

We thank Bruce Rosengarten and Wayne Cawthorne for helpful discussions; WEHI Bioservices, WEHI Histology, the WEHI Monoclonal Antibody Facility for technical assistance; Gary Kasof and Cell Signaling Technology for generously sharing antibodies; Professor Marc Pellegrini for the *Casp8^fl/fl^* strain; and Tina Cardamone and Dr John Finnie from Phenomics Australia Histopathology and Slide Scanning Service for their assistance with phenotyping the *Ripk3^D142N/D142N^* mouse strain. The generation of *Ripk3^D143N/D143N^* mice used in this study was supported by Phenomics Australia and the Australian Government through the National Collaborative Research Infrastructure Strategy (NCRIS) program.

## FUNDING

This work was supported by National Health and Medical Research Council of Australia grants (1172929 and 2034104 to JMM, 2008652 to EDH, 2002965 to ALS, 2034044 to CRH; IRIISS 9000719), and by the Victorian State Government Operational Infrastructure Support scheme. TLP is supported by a Viertel Senior Medical Research Fellowship. JMM received research funding from Anaxis Pharma Pty Ltd and AH was funded by Anaxis Pharma Pty Ltd. We acknowledge scholarship support for SC (Australian Government Research Training Program Stipend Scholarship, WEHI Handman PhD Scholarship).

## AUTHOR CONTRIBUTIONS

Conceptualization: SC, EDH, ALS, JMM. Investigation: SC, KMP, AP, JAR, CRH, SNY, SEG, AH, CH, JMH, AJK, ALS; Methodology: SC, KMP, AJK, TLP, ALS. Resources: TLP, EDH, ALS, JMM. Supervision: EDH, ALS, JMM. Funding acquisition: EDH, ALS, JMM. Writing: SC, ALS and JMM co-wrote the paper with input from authors.

## COMPETING INTERESTS

KMP, CRH, SNY, AH, JMH, ALS and JMM contribute to or have contributed to a project developing necroptosis inhibitors in collaboration with Anaxis Pharma. TLP is cofounder and shareholder in Nelcanen Therapeutics. The other authors declare no competing interests.

## ETHICS DECLARATION

All experiments were approved by the WEHI Animal Ethics Committee following the Prevention of Cruelty to Animals Act (1996) and the Australian National Health and Medical Research Council Code of Practice for the Care and Use of Animals for Scientific Purposes (1997).

