## Supplemental Figures 1-5, Supplemental Tables 1-2 for "The kinase domain of RIPK3 tunes its scaffolding functions"

**Supplementary Figure 1. The RIPK3 inhibitor, GSK′843, induces apoptosis by driving RIPK3 into cytoplasmic puncta.** (**A**) 10ng/mL doxycycline (DOX) was used to reconstitute RIPK3 variant expression in *Ripk3*$^{-/-}$ MDF cells to emulate endogenous expression levels. Representative of n=4 for RIPK3$^{WT}$, RIPK3$^{K51A}$, RIPK3$^{D143N}$, and n=1 for RIPK3$^{D161N}$. (**B**) GSK′843 induced formation of cytoplasmic RIPK3 puncta in *Ripk3*$^{-/-}$ MDF cells reconstituted with wild-type RIPK3 in the presence of the Caspase inhibitor, IDN-6556. (**C**) GSK′843 increases the thermal stability of endogenous RIPK3 in wild-type MDF cells, illustrating on-target binding, in immunoblots of post-treatment cell lysates. GAPDH is included as a loading control.

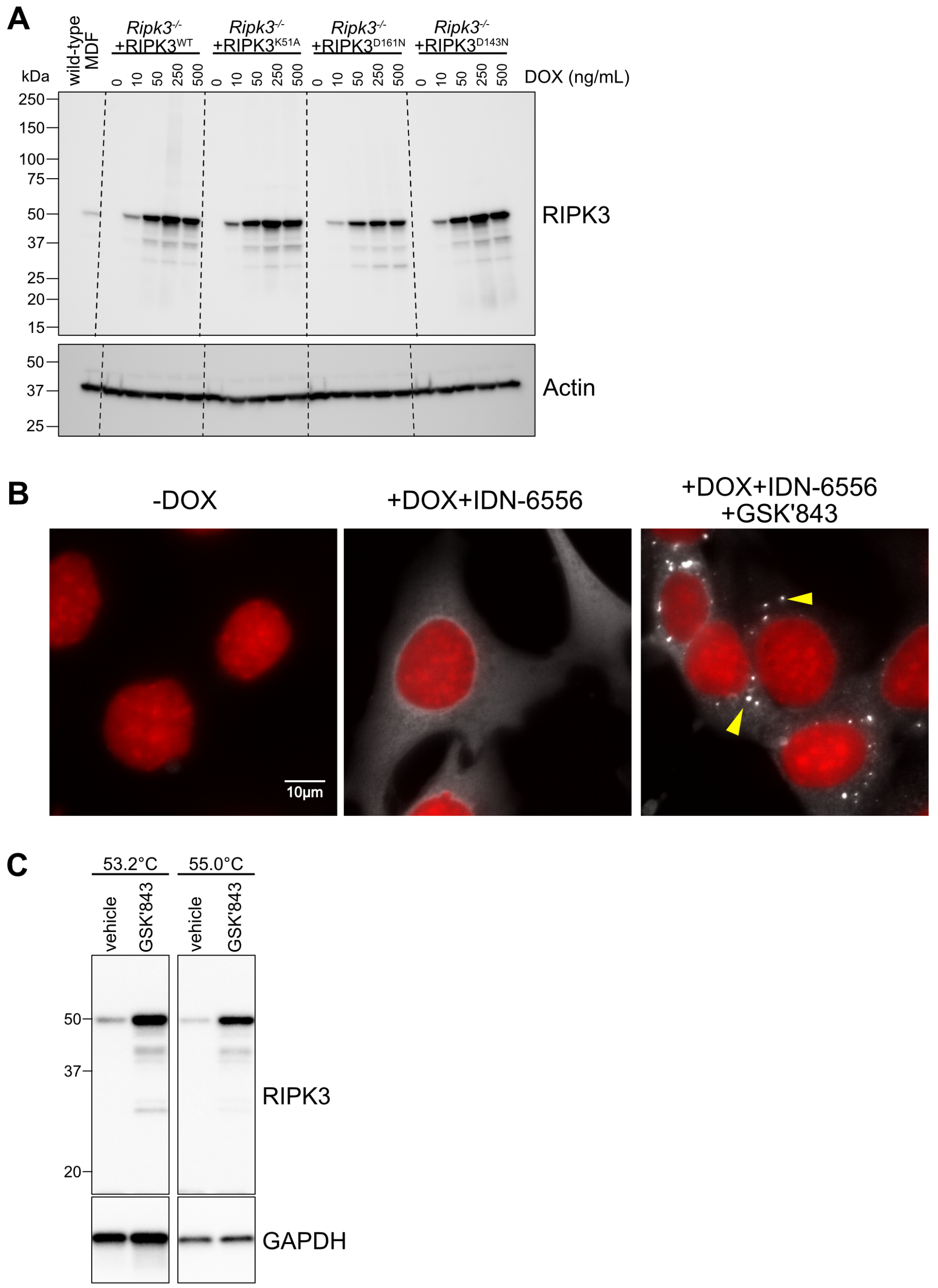

**Supplementary Figure 2. No developmental defects in unchallenged *Ripk3*$^{D143N}$ compared to wildtype littermates up to one year of age.** **A** H&E stain of **A** No macroscopic defects in *Ripk3*$^{D143N/D143N}$ mice at 7-10 months of age compared to *Ripk3*$^{WT/WT}$ littermates. Blinded comprehensive examinations performed by veterinary pathologists indicated that at the time of necropsy, the mice appeared well nourished, well groomed, active/curious (rearing on hind legs) and healthy with normal movement and gait. There were no observable dermal lesions, typical oral features, unremarkable dentition and no nasal/ocular discharges. The coat appeared well groomed and smooth. No observable abnormalities of the limbs, paws and digits. The gastrointestinal tract contained ample ingesta and the thoracic and abdominal viscera showed no macroscopic abnormalities. $n$=3 per genotype for all organs except testes $n$=2 per genotype. Scale bar represents 1000 µm for brain images, 100 µm for testis images and 500 µm for eye, ileum and spinal cord images. **B** Weight of *Ripk3*$^{D143N}$ and wildtype littermates. Wildtype $n$= 23, and *Ripk3*$^{D143N}$ $n$=13. **C** Red blood cell count of *Ripk3*$^{D143N}$ and wildtype littermates up to one year old. **D** Neutrophil count of *Ripk3*$^{D143N}$ and wildtype littermates up to 13 months of age. **E** Lymphocyte count of *Ripk3*$^{D143N}$ and wildtype littermates up to 13 months of age. **F** Monocyte count of *Ripk3*$^{D143N}$ and wildtype littermates up to 13 months of age. **G** Eosinophil count of *Ripk3*$^{D143N}$ and wildtype littermates up to 13 months of age. For panels C-G, Wildtype $n$= 48, and *Ripk3*$^{D143N}$ $n$=35.

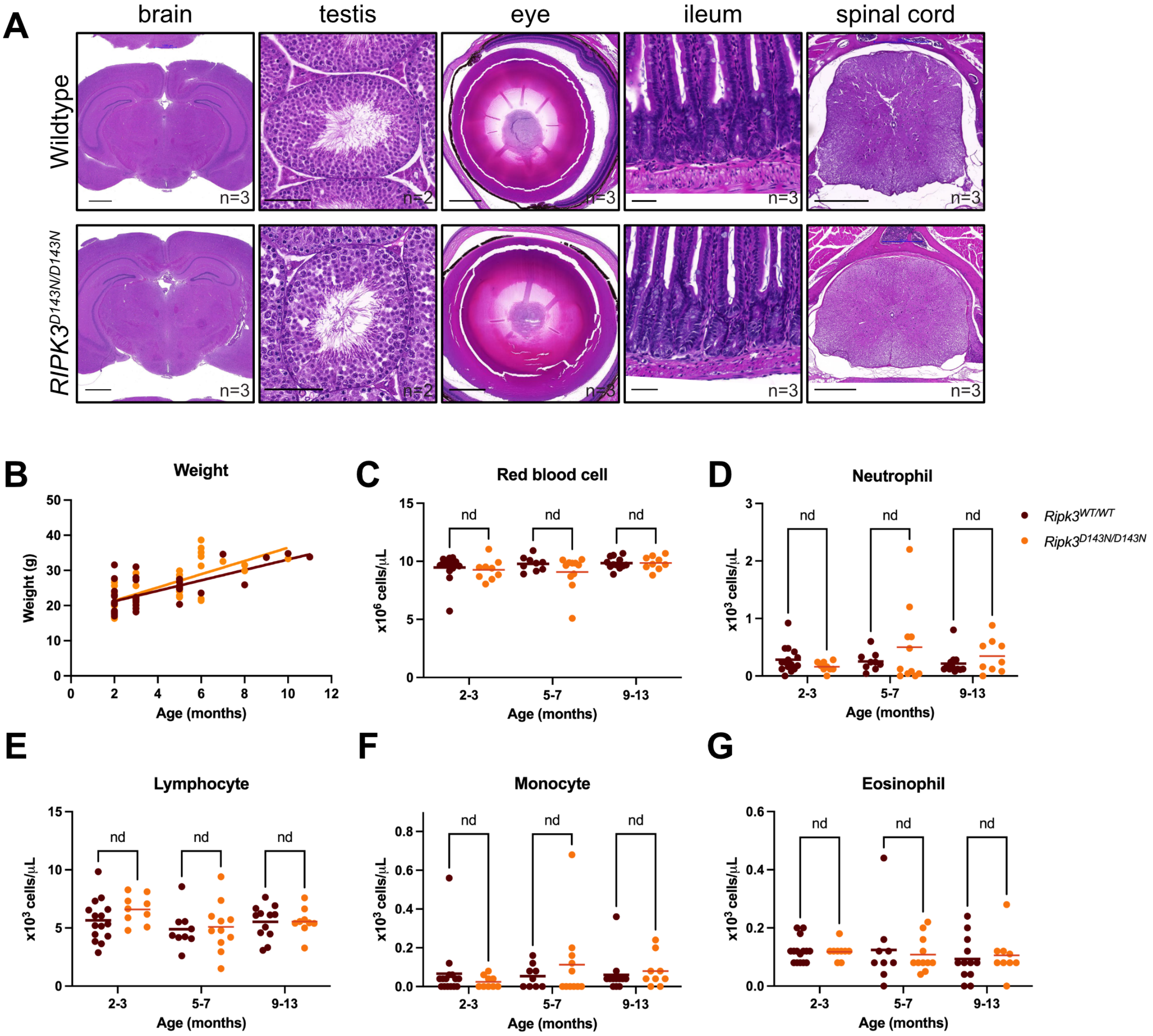

**Supplementary Figure 3. RIPK3$^{D143N}$ was expressed and distributed comparably to RIPK3$^{WT}$ across tissues, and did not perturb Caspase-8, RIPK1 and Cleaved Caspase-3 abundance, in the absence of challenge.** Immunohistochemical staining of Caspase-8, RIPK1, RIPK3 and Cleaved Caspase-3 in organs from wild-type and *Ripk3$^{D143N}$* knock-in mice. Scale bar represents 500 µm for spleen, kidney, liver, heart, and 50 µm for small intestine, large intestine.

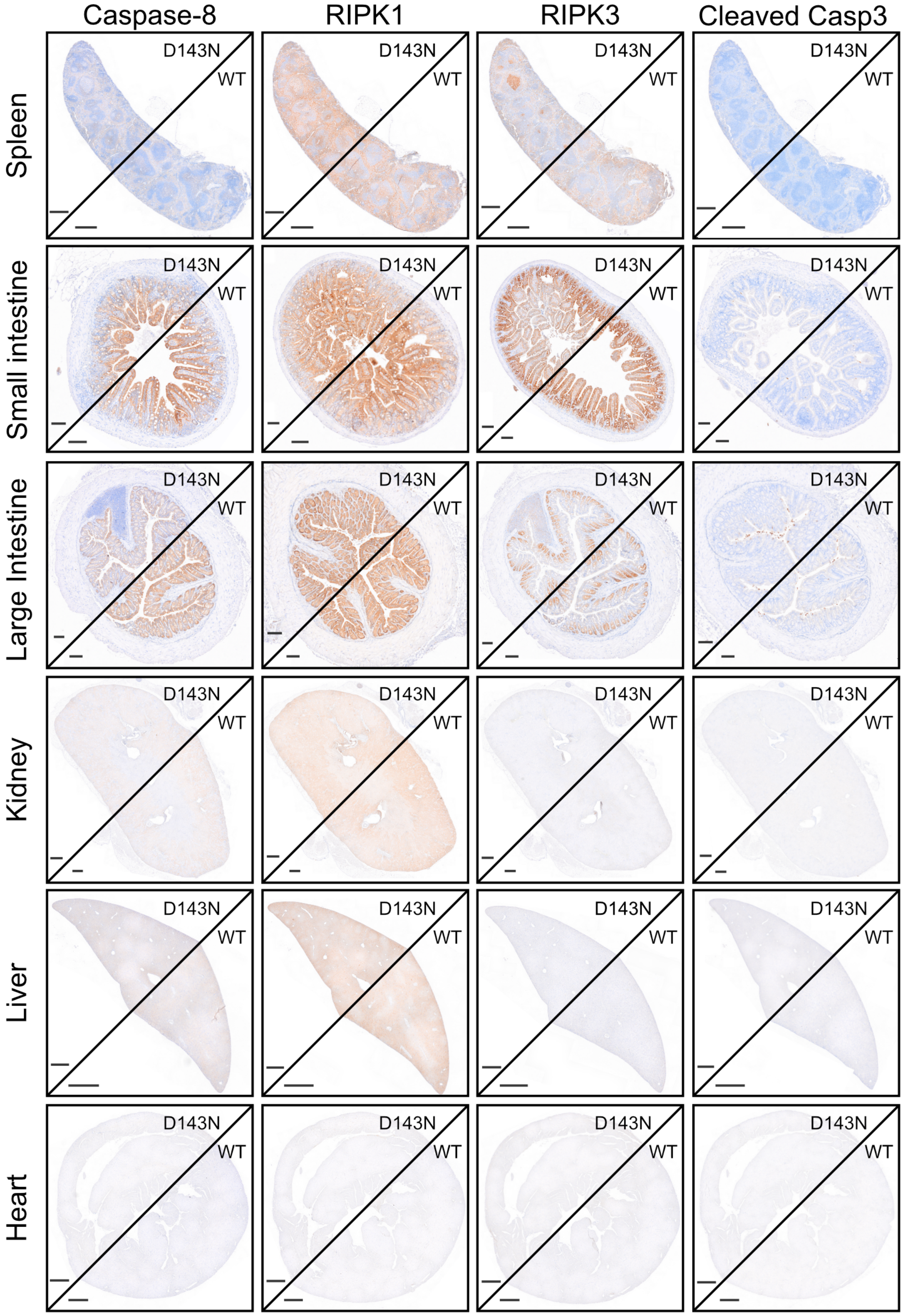

**Supplementary Figure 4. Villin.Cre1000 is expressed in the gut epithelium with modest expression in the liver. A** ROSA protein expression in wild-type and *ROSA26 Ai9* Villin.Cre1000 (ROSA Villin.Cre1000) mouse organs. Images representative of *n*=1 for wildtype and *n*=5 for ROSA Villin.Cre1000 mice. Scale bar represents 100 µm. **B** Immunohistochemistry confirms that crossing Villin.Cre1000 with $Casp8^{fl/fl}$ mice eliminated expression of Caspase-8 in ileum. Images representative of *n*=3 wild-type, *n*=5 $Casp8^{IEC}$, *n*=4 for both $Casp8^{IEC}Mlkl^{IEC}$ and $Casp8^{IEC}Ripk3^{D143N}$ mice. Scale bar represents 20 µm.

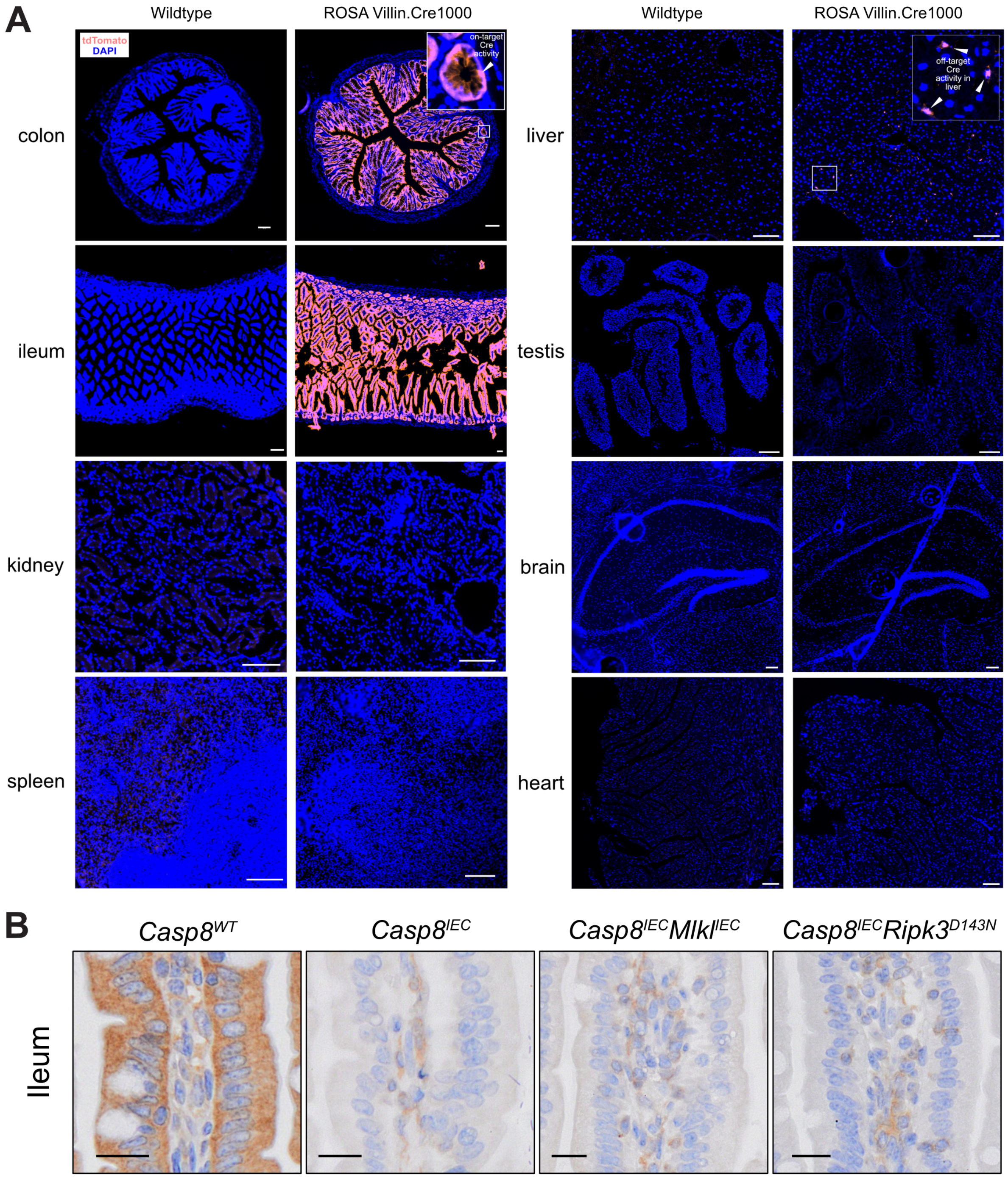

**Supplementary Figure 5. Paneth cell loss in *Casp8$^{IEC}$* mice is restored by MLKL deletion in IEC cells or systemic RIPK3$^{D142N}$ mutation.** Paneth cells were detected by immunohistochemical staining of MPTX2 in the ileum (**A**) and enumerated (**B**). *n*=3 wild-type and *n*=4 for each of *Casp8$^{IEC}$*, *Casp8$^{IEC}$Mlkl$^{IEC}$* and *Casp8$^{IEC}$Ripk3$^{D143N}$* mice. Bars indicate group mean ±SD. n.s. indicates p>0.05 by ordinary one-way ANOVA with Tukey's correction. (**C**) *Casp8$^{IEC}$* deletion did not cause significant mortality. Survival curve of wild-type and *Casp8$^{IEC}$* mice littermates up to 6 months of age analyzed with simple Survival Analysis Kaplan-Meier Log-rank Mantel-Cox test. *n*=6 for wild-type (*Casp8$^{WT}$*) and *n*=19 *Casp8$^{IEC}$* mice. (**D**) *Casp8$^{IEC}$* mice gain weight normally up to 6 months of age comparable to wildtype littermates. Mean±SEM graphed, male (M) wild-type *n*=6, male *Casp8$^{IEC}$* *n*=6, female (F) wild-type *n*=7, and female *Casp8$^{IEC}$* *n*=6.

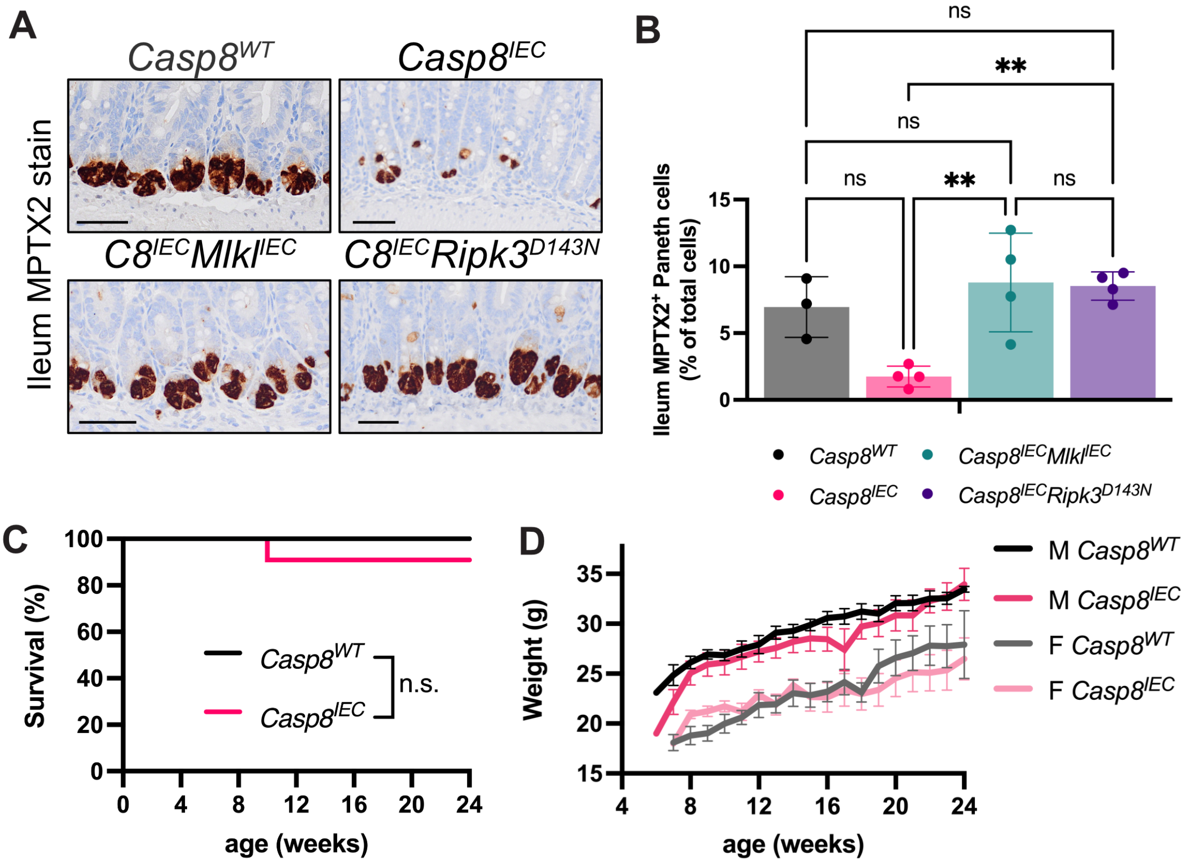

**SUPPLEMENTAL TABLES**

**Supplementary Table 1. Immunohistochemistry protocols.**

| | **Procedures in Lysozyme stain protocol on Dako Omnis** | **Time** |
|---|---|---|
| 1 | Deparaffinization onboard<br>Phase 1: Clearify Clearing Agent<br>Phase 2: DI water | |
| 2 | Heat-induced epitope retrieval<br>EnVision FLEX TRS, High pH retrieval buffer onboard | 30 min |
| 3 | Primary antibody<br>1:1500 Lysozyme | 1 hr |
| 4 | Endogenous enzyme blocking<br>Dako REAL Peroxidase-blocking reagent | 5 min |
| 5 | Secondary labelled Polymer<br>Anti-Rabbit HRP polymer | 30 min |
| 6 | Chromogen-substrate<br>DAB | 8 min |
| 7 | Counterstain<br>In-house made Mayer’s Hematoxylin, followed by Bluing reagent | 1 min each |
| 8 | Dehydrate and coverslip | |
| | Procedures in MPTX2 stain protocol on Dako Omnis | Time |
| 1 | Deparaffinization onboard<br>Phase 1: Clearify Clearing Agent<br>Phase 2: DI water | |
| 2 | Heat-induced epitope retrieval<br>EnVision FLEX TRS, Low pH retrieval buffer onboard | 30 min |
| 3 | Primary antibody<br>1:1000 MPTX2 (Abcam, Cat#EPR20920-19) | 1 hr |
| 4 | Endogenous enzyme blocking<br>Dako REAL Peroxidase-blocking reagent | 4 min |
| 5 | Secondary labelled Polymer<br>Anti-Rabbit HRP polymer | 30 min |
| 6 | Chromogen-substrate<br>DAB | 10 min |
| 7 | Counterstain<br>In-house made Mayer’s Hematoxylin, followed by Bluing reagent | 1 min each |
| 8 | Dehydrate and coverslip | |
| | Procedures in PAS stain protocol | Time |

| | | |
|---|---|---|
| 1 | Dewax sections and bring to water in usual manner | |
| 2 | 1% Aqueous Periodic Acid | 15 min |
| 3 | Wash in distilled water | 1 min |
| 4 | Schiff's Reagent (Sigma-Aldrich,1090330500) | 20 min |
| 5 | Wash in water | 5 min |
| 6 | Mayer's Hematoxylin | 1 min |
| 7 | Wash in water | 1 min |
| 8 | Scott's tap water (Leica, 3802915) | 1 min |
| 9 | Wash in water | 1 min |
| 10 | Dehydrate, clear and mount | |

**Supplementary Table 2. Quantification scripts.**

| **PAS$^+$ cell detection** |
|---|
| runPlugin('qupath.imagej.detect.cells.WatershedCellDetection', '{"detectionImageBrightfield":"Hematoxylin OD","requestedPixelSizeMicrons":0.2739,"backgroundRadiusMicrons":0.0,"backgroundByReconstruction":true,"medianRadiusMicrons":0.0,"sigmaMicrons":1.5,"minAreaMicrons":10.0,"maxAreaMicrons":400.0,"threshold":0.05,"maxBackground":2.0,"watershedPostProcess":true,"excludeDAB":false,"cellExpansionMicrons":5.0,"includeNuclei":true,"smoothBoundaries":true,"makeMeasurements":true}')// |
| **Total cell count** |
| runPlugin('qupath.imagej.detect.cells.WatershedCellDetection', '{"detectionImageBrightfield":"Hematoxylin OD","requestedPixelSizeMicrons":0.2739,"backgroundRadiusMicrons":10.0,"backgroundByReconstruction":true,"medianRadiusMicrons":0.0,"sigmaMicrons":1.5,"minAreaMicrons":10.0,"maxAreaMicrons":200.0,"threshold":0.05,"maxBackground":2.0,"watershedPostProcess":true,"excludeDAB":false,"cellExpansionMicrons":5.0,"includeNuclei":true,"smoothBoundaries":true,"makeMeasurements":true}') |
| **Lysozyme$^+$ cell detection and total cell count** |
| runPlugin('qupath.imagej.detect.cells.PositiveCellDetection', '{"detectionImageBrightfield":"Hematoxylin OD","requestedPixelSizeMicrons":0.2739,"backgroundRadiusMicrons":0.0,"backgroundByReconstruction":true,"medianRadiusMicrons":0.0,"sigmaMicrons":1.5,"minAreaMicrons":20.0,"maxAreaMicrons":700.0,"threshold":0.14,"maxBackground":10.0,"watershedPostProcess":true,"excludeDAB":false,"cellExpansionMicrons":5.0,"includeNuclei":true,"smoothBoundaries":true,"makeMeasurements":true,"thresholdCompartment":"Cell: DAB OD mean","thresholdPositive1":0.08,"thresholdPositive2":0.4,"thresholdPositive3":0.6000000000000001,"singleThreshold":false}') |
